# Meal timing is a critical factor for maintenance of gut homeostasis around the clock

**DOI:** 10.1101/2025.08.12.669757

**Authors:** Felicity K Hunter, Polly Downton, Andrea Luengas-Martinez, Suzanna H Dickson, Jafar Cain, Kathryn J Else, Matthew R Hepworth, Julie E Gibbs

## Abstract

Gut function exhibits 24 hour (circadian) rhythmicity, in part driven by intrinsic clocks within intestinal epithelial cells (IECs). The gut microbiome also demonstrates circadian rhythms in composition and function, important for maintenance of metabolic, immune and gut health. Here, we determined the influence of feeding behaviour on the colonic circadian landscape using an interval feeding paradigm, whereby food intake was partitioned equally across the 24h day. RNAseq analysis revealed that the IEC intrinsic clock persists in the absence of diurnal feeding rhythms, however a subset of key transcripts lose rhythmicity, demonstrating that cell extrinsic temporal cues contribute significantly to maintenance of the rhythmic gut transcriptome. Furthermore, interval fed mice demonstrated striking loss of rhythms in secretory IgA, a critical regulator of the temporal landscape of the gut microbiome. In keeping, rhythmicity within the microbiota and microbial derived short chain fatty acids was significantly diminished. This work highlights the importance of daily rhythms in feeding behaviour for maintenance of rhythmic processes within the gut, with implications for metabolic and immune health.

## Introduction

The circadian clock is an intrinsic timer which regulates biological processes to align with the 24 hour (24h) day. In mammals, light information is conveyed to the central clock within the hypothalamus via the retino-hypothalamic tract, thereby tuning it to daily light-dark cycles^(1)^. In turn, the central clock coordinates peripheral clocks, found in most cells and tissues around the body, via non-photic *zeitgebers* (meaning time givers) including neuronal signals, hormones, temperature and food intake^(2)^. At the molecular level, a handful of core clock genes/proteins (*Bmal1*, *Period1/2*, *Clock*, *Cryptochrome 1/2*, *Nr1d1/2* and *Ror*) form transcriptional translational feedback loops which drive 24h rhythms in clock-controlled genes to facilitate tissue specific circadian programmes of rhythmic biology^(3, 4)^. While co- ordination of circadian rhythmicity via the central clock has been extensively studied, recent advances suggest peripheral cues, including those from the diet, also act to entrain rhythmic biology.

Gastrointestinal (GI) functions including colonic motility, permeability, localised hormone secretion and immune function exhibit 24h rhythmicity^(5)^ ^(6, 7)^. Our earlier work^(8)^, and that of others^(6, 9–12)^, demonstrates that over 50% of the colonic transcriptome is rhythmic in *ad libitum* fed mice. At a cellular level, intestinal epithelial cells (IECs) house a robust intrinsic clock. Cell-specific ablation of the IEC clock (via targeted deletion of the core clock gene *Bmal1*) causes widespread loss of 24h rhythmicity within the colonic transcriptome, although a significant proportion of rhythmic transcripts remain unchanged^(13)^. This is suggestive of either additional timing cells within the colon and/or a role for extrinsic rhythmic signals for driving tissue rhythms independent of the IEC molecular clock. IECs form a single lining layer along the gut and play an important role in maintaining gut homeostasis^(14)^. Critically, IECs along with underlying immune responses (including secretory Immunoglobulin A (IgA)), segregate the gut microbiota from the host, and facilitate crosstalk between the microbiome and gut resident immune cells. The gut microbiota exhibits 24h oscillations in composition, biophysical localisation within the intestine and metabolic outputs^(9, 15–19)^. This daily rhythmicity in the microbiome is important for maintenance of metabolic and immune health^(20–22)^.

Feeding behaviour is highly influential over peripheral clocks^(23)^. Bouts of feeding usually occur during the active phase of the 24h cycle, daytime for humans and night-time for rodents. Studies in rodents demonstrate that under reverse feeding cycles, where food availability is restricted to the daytime (and thus mis-aligned to the active phase), highly metabolic organs such as the liver exhibit altered diurnal rhythms in expression of core clock genes^(24, 25)^ and other hepatic transcripts^(26)^, which re-align with food availability. Molecular mechanisms by which food intake synchronises clockwork machinery are not fully understood, but feeding derived signals originating from the host^(27)(28)^ and the gut microbiota^(29)^ can re-set the clock.

Here we sought to address the influence of feeding behaviour on the circadian landscape of the colon using an interval feeding paradigm whereby food intake was partitioned equally across the 24h day. We focused on the IEC, a key timer cell within the colon and interface between the host and gut microbiota. In the absence of diurnal rhythms in feeding behaviour, the core clock mechanism persisted within IECs. However, there was significant re-organisation of transcriptional rhythmicity within the gut, and loss of rhythmicity in secretory IgA, beneficial commensals and short chain fatty acids. Together these data highlight the importance of feeding behaviour for synchronising the gut clock and sustaining rhythmicity, demonstrating that meal timing is a critical factor for maintenance of gut homeostasis.

## Results

### Interval feeding uncouples rhythms of food intake from rhythmic behaviours and circadian outputs

To determine the influence of diurnal feeding behaviour on the IEC clock and the circadian transcriptome we established an interval feeding regimen. Mice were provided with 8 small meals a day at 3 hourly intervals for 16 days (**Figure 1A**). This resulted in an equal spread of food consumption across each mealtime (**Figure 1B).** In contrast, *ad libitum* fed mice consumed the majority of their food at night, with approximately 21% of their food eaten during the light period, as expected (**Figure 1B).** During the early acclimatisation period (days 1 and 2), daily food intake was reduced under interval feeding (**Supplementary Figure 1A**). However, mice quickly adapted to the regimen, consuming similar amounts of food each day, and consistently consuming 12-13% of total daily intake at each meal. Whilst *ad libitum* fed mice gained weight (∼8%) over the experimental period, interval fed mice maintained their starting weight (**Supplementary Figure 1B, C**). Daily activity and metabolic parameters were analysed using the PhenoMaster system. Activity transiently spiked at mealtimes both in interval fed and *ad libitum* fed mice (co-housed within the same light-tight cabinet) (**Figure 1C**). Nevertheless, under both feeding regimens mice were predominantly active during the night, demonstrating maintenance of nocturnal behaviour. Similarly drinking activity was greater in the night versus day (**Figure 1D**). Oxygen consumption (VO_2_), carbon dioxide production (VCO_2_) and heat production transiently spiked during mealtimes in interval fed mice, but in keeping with *ad libitum* fed controls, increased at the onset of the dark phase (**Supplementary Figure 1D**-F). Respiratory exchange ratio (RER), dipped during the onset of night in interval fed mice, indicating a shift in fuel source from carbohydrates towards fatty acids, but increased transiently when meals were consumed (reflective of maintained nocturnal activity, but reduced food intake during the dark period) (**Figure 1E**). Finally, circulating levels of the circadian hormone corticosterone remained rhythmic with a peak before the onset of the dark phase under both feeding regimens (**Figure 1F**). To summarise, mice maintained 24h rhythms in behavioural activity and corticosterone levels under interval feeding, further supporting the effectiveness of an interval feeding protocol to uncouple rhythms of food intake from other rhythmic behaviours and circadian outputs^(30–35)^. These data shed new light on the influence of interval feeding over metabolic parameters, revealing that overall metabolic rate was not perturbed.

**Figure 1:**
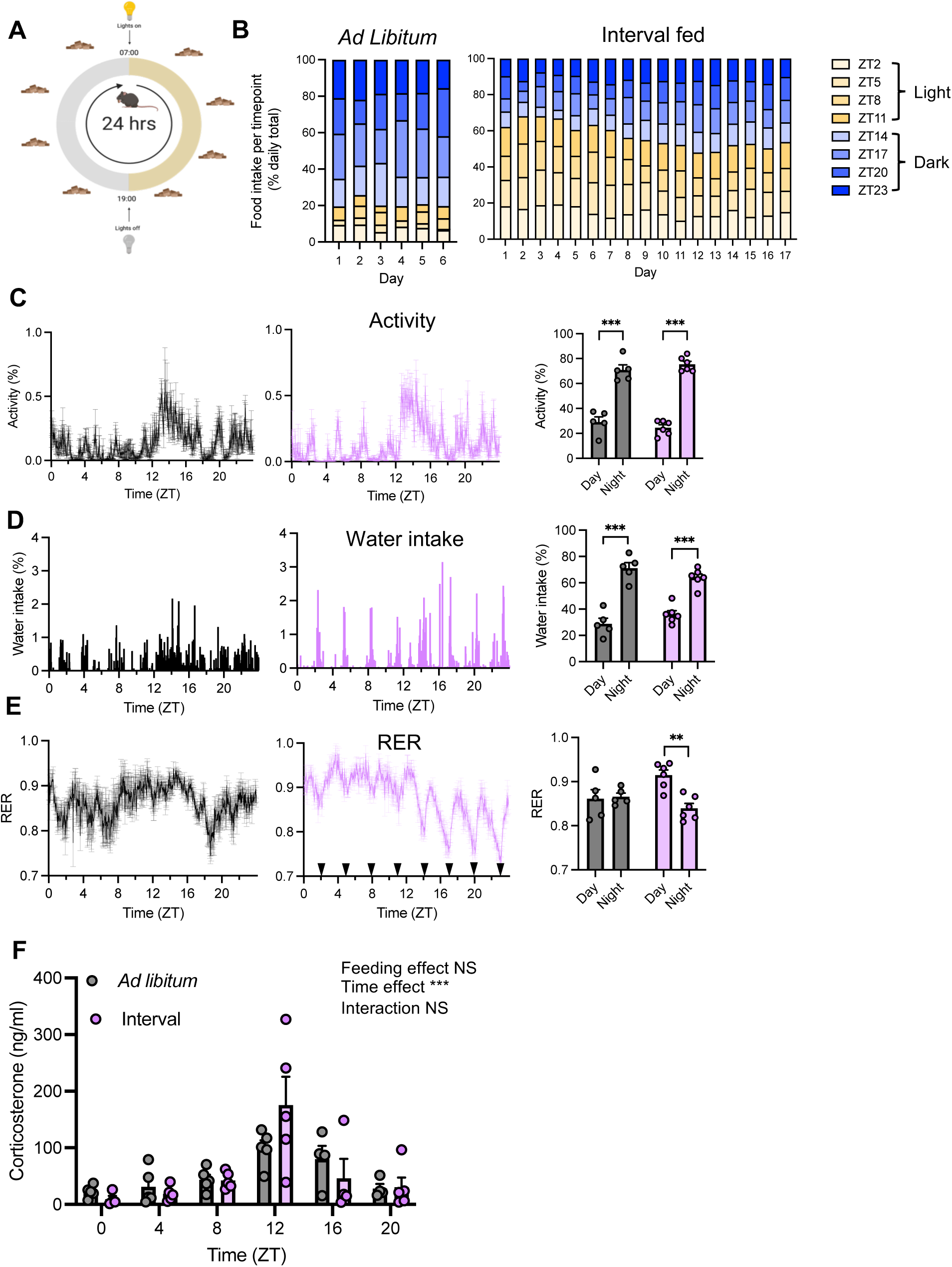
Effects of interval feeding on food intake and rhythmic physiology. (A) Schematic illustrating the interval feeding schedule. Mice were maintained under a 12:12 light dark cycle and provided access to food for a short window of time every 3 hours. (B) Quantification of food intake at every mealtime. Food intake per cage (5 animals per cage, averaged over 3 cages (*ad libitum* fed) or 6 cages (interval fed)) and expressed as a percentage of the days’ total intake. Phenomaster cages were utilised to measure (in singly housed mice): (C) activity; (D) water intake and (E) respiratory exchange rate (RER) under *ad libitum* feeding (black) and interval feeding (purple). Line plots show group average (n=6 interval fed, n=5 *ad libitum* fed), meal times denoted by triangles. Bar charts quantify partitioning into day (12h lights on) and night (12h lights off), 2 way ANOVA and post-hoc Tukey’s multiple comparisons test. (F) Circulating corticosterone levels in *ad libitum* fed mice (n=4-5/time point) and interval fed mice (n=4-5/time point), 2 way ANOVA. *See also Supplementary* Figure 1.

### Interval feeding drives temporal reorganisation of the IEC transcriptome, despite persistence of the cellular molecular clock

We utilised the interval feeding paradigm to reveal how the IEC circadian transcriptome responds to loss of diurnal feeding behaviour. IECs (∼80% purity, **Supplementary Figure 2A**) were harvested across the 24h day from interval fed mice and *ad libitum* controls for RNAseq analysis. Interval feeding drove substantial reorganisation of 24h rhythms in gene expression (**Figure 2A**). Whilst 3662 transcripts were similarly rhythmic under both regimens (same, **Supplementary Figure 2A**), a proportion (217 transcripts) demonstrated a change in rhythmicity (altered amplitude and/or phase, **Figure 2B**), whilst others gained (418 transcripts) or lost (1237 transcripts) rhythmicity. Of note, approximately 25% of the IEC circadian transcriptome lost rhythmicity under interval feeding. Loss and gain of rhythmicity were not owing to changes in overall expression levels (**Supplementary Figure 2C**), indicating that transcript oscillation is regulated independently of magnitude of expression. Furthermore, analysis of the two feeding regimens (independent of sample collection time) revealed no differentially expressed genes (DESeq2 analysis, padj<0.1, data not shown). This indicates that the temporal dynamics of transcription, but not the transcriptional programmes themselves, are altered by interval feeding.

**Figure 2:**
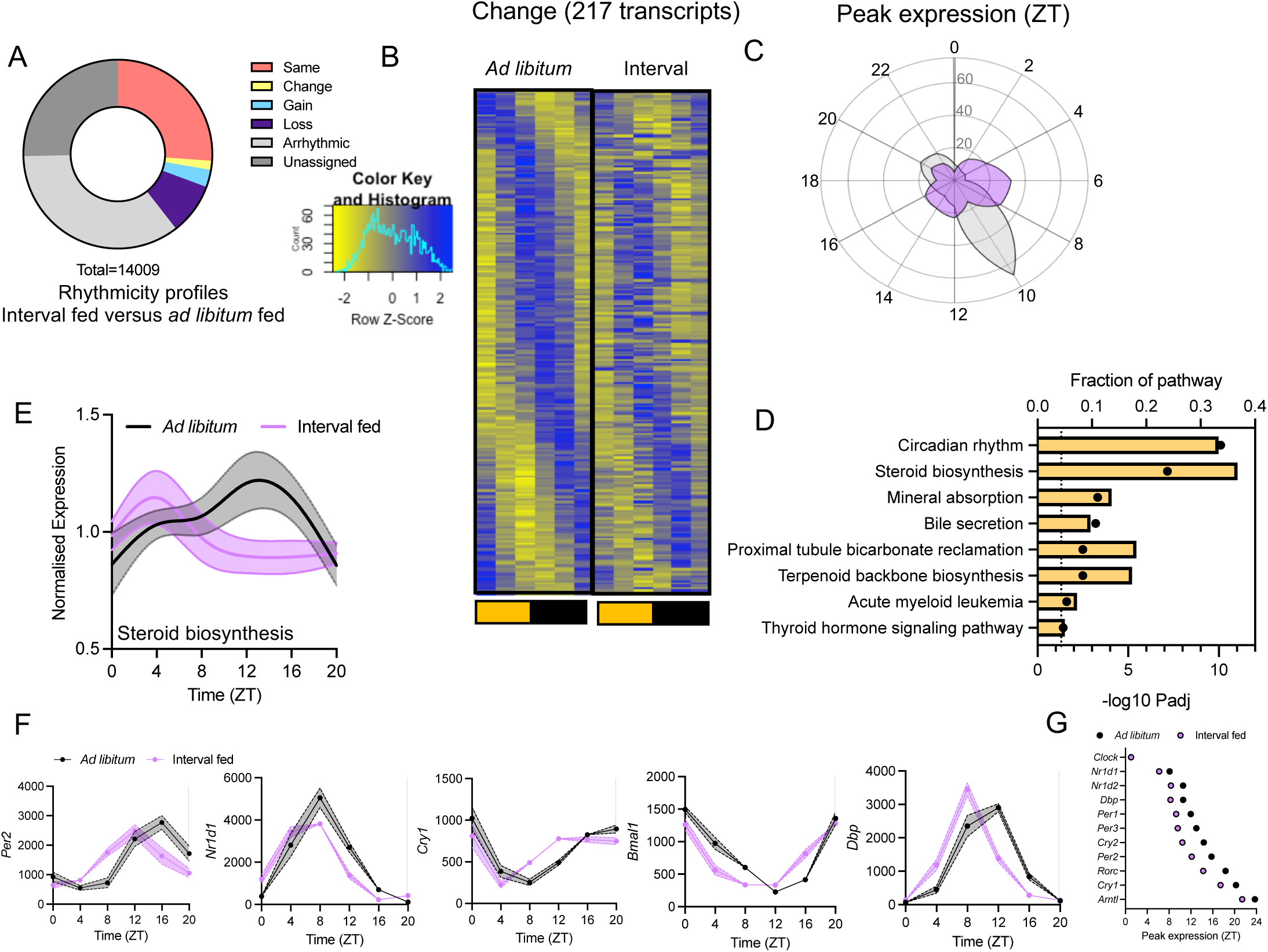
The IEC clock persists under interval feeding. (A) Proportion of the IEC transcriptome showing 24h rhythms under both *ad libitum* and interval feeding (“same” or “change”); under *ad libitum* feeding only (“loss”); or under interval feeding only (“gain”). 24h rhythmicity assessed using CompareRhythms. (B) Transcripts demonstrating changed rhythmicity under interval feeding, n=4-5/timepoint, 6 time points per condition. (C) Temporal distribution of the peak expression of transcripts demonstrating changed rhythmicity, grey = *ad libitum* and purple = interval fed. (D) Pathway analysis (mouse KEGG 2019) of transcripts assigned a change in rhythmicity. Bars quantify fraction of the pathway represented, and dots represent statistical significance (dotted line marks Padj=0.05). (E) Spline plot of normalised expression of all genes in the “Steroid biosynthesis” pathway which appeared in our data (Mmu 00100; 17 genes), error bars represent 95% CIs around the mean. (F) Expression of clock genes (normalized expression) over time which were assigned changed rhythmicity, n=4-5/time point. (G) Phasing of clock gene expression within the IECs (as determined by CompareRhythms) under *ad libitum* or interval feeding. *See also Supplementary* Figures 2 and 3.

Examination of the changed transcript subset revealed that in *ad libitum* fed mice, expression peaked predominantly at transition from light to dark (prior to the onset of activity and increased food intake). Under interval feeding this phasing was re-distributed across the 24h day (**Figure 2C**). Further analysis mapped these changed transcripts onto pathways including “circadian rhythms”, “steroid biosynthesis” (including *Fdft1*, *Sqle* and *Cyp51*), “bile secretion” (including *Ldlr* and *Hmgcr*) and “mineral absorption” (including *Mt1/2* and *Atp1a1*) (**Figures 2D and E, Supplementary Figure 2D**). Importantly, the majority of core clock genes fell into the change group with *Per1/2/3*, *Cry1/2, Arntl (Bmal1)*, *Dbp* and *Rorc* exhibiting robust 24h rhythms under interval feeding. However, these rhythms were phase advanced by 2–4h (with the exception of *Clock*), and thus misaligned with observed activity and hormonal rhythms (**Figure 2F and G** and **Supplementary Figure 2E**). Similar shifts in clock gene phase were observed in the liver (**Supplemental Figure 3**) comparable to findings by others employing similar feeding regimens^(30)^. Thus, the IEC clock persists in the absence of diurnal rhythms in feeding behaviour, but rhythmic feeding provides temporal cues which synchronise this timer to the light dark cycle.

### Diurnal feeding behaviour delivers rhythmic cues important for maintenance of the IEC circadian transcriptome

The observed loss and gain of rhythmicity in subsets of transcripts under interval feeding, despite the intact clockwork machinery, demonstrates dependence on rhythmic cues delivered by diurnal feeding behaviour. This observation provides new insight into the relative importance of extrinsic temporal cues and intrinsic clocks in the maintenance of a rhythmic gut transcriptome. Transcripts which gained rhythmicity under interval feeding tended to peak during the middle of the day (ZT6-8) (**Figure 3A and B**) and could be mapped to a small number of pathways including “glycerophospholipid metabolism” (including *Pla2g12a* and *Lpin2*) and “ether lipid metabolism” (including *Plpp3* and *Pld2*) (**Figure 3C and D**).

**Figure 3:**
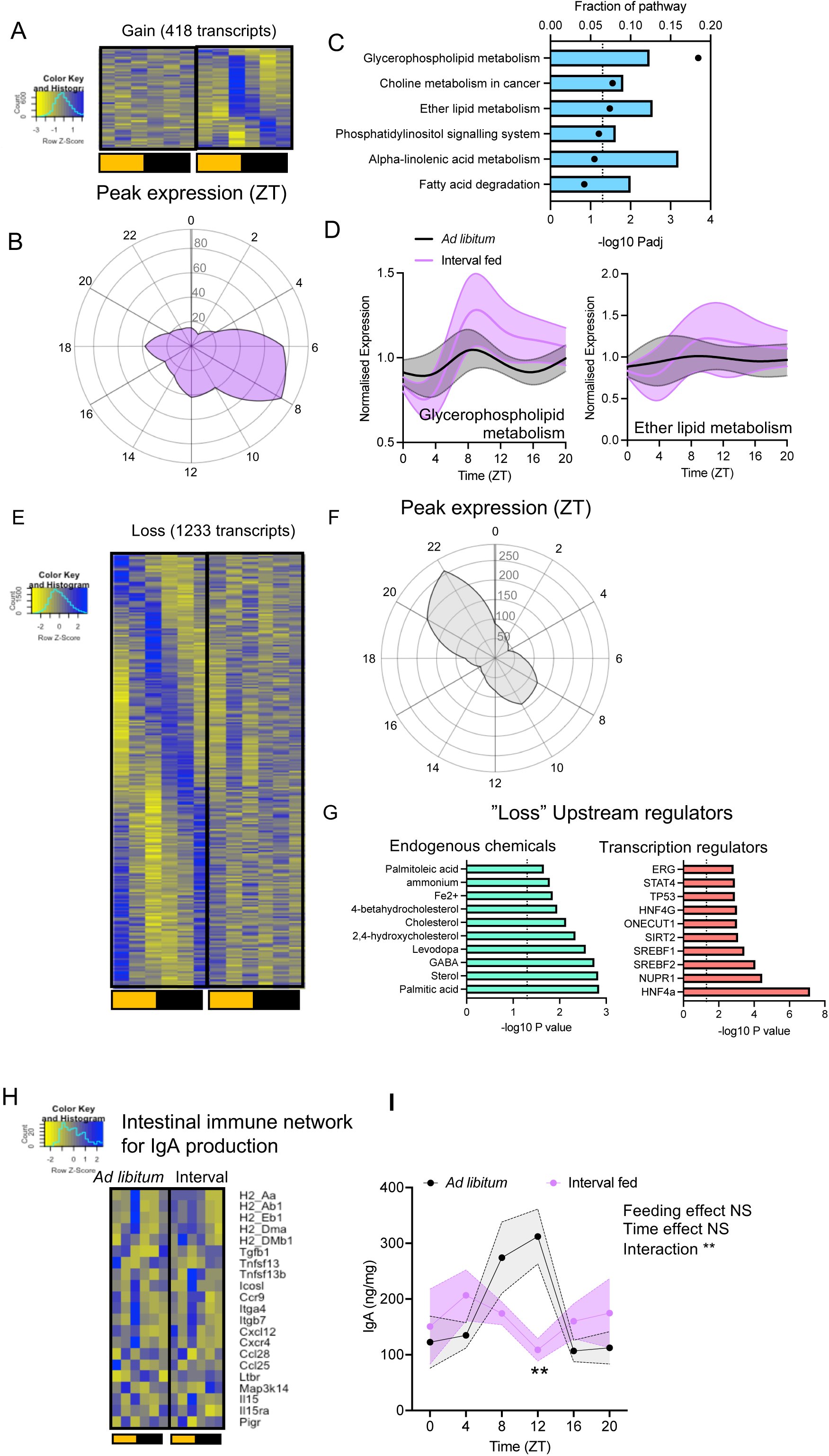
Interval feeding drives significant temporal re-organisation of the IEC transcriptome. (A) 418 transcripts showed gain of 24h rhythmicity under interval feeding. (B) Temporal distribution of the peak expression of transcripts which were arrhythmic in *ad libitum* fed mice but gained 24h rhythmicity under interval feeding. (C) Pathway analysis (mouse KEGG 2019) of transcripts assigned a gain in rhythmicity. Bars quantify fraction of the pathway represented, and dots represent statistical significance (dotted line marks Padj=0.05). (D) Spline plots of mean normalized expression of all genes in the denoted pathways which appeared in the data and were in the gain category, error bars represent 95% CIs around the mean. Glycerophospholipid metabolism (Mmu_00564, 12 genes) and Ether lipid metabolism (Mmu_00565, 6 genes). (E) 1233 transcripts showed loss of 24h rhythmicity under interval feeding. (F) Temporal distribution of the peak expression of transcripts which were rhythmic in *ad libitum* fed mice but arrhythmic under interval feeding. (G) Upstream regulator analysis of IEC transcripts which lost rhythmicity under interval feeding. The top 10 regulators in the categories “endogenous chemicals” and “transcription regulators” and their significance are shown. (H) Normalised expression across time of genes within the IEC RNAseq data which were in the KEGG mouse 2019 pathway “intestinal immune network for IgA production” (mmu_04672; 21 genes), n=4-5/timepoint. (I) Fecal IgA across time in *ad libitum* and interval fed animals, n=3-5/time point. 2 way ANOVA and post hoc Sidak’s multiple comparisons tests between feeding regimens at each time point. *See also Supplementary* Figure 4.

Transcripts which lost rhythmicity tended to peak under *ad libitum* conditions at either the end of the night (ZT20-22) or end of the day (ZT8-10) (**Figures 3E and F**). Enrichment analysis did not yield any significant common biological pathways (using adj.P<0.05), however less stringent analysis (using P<0.05**, Supplementary Figure 4A**) did identify pathways including “citrate cycle” and “lysine degradation”. Analysis of upstream regulators (IPA analysis, **Figure 3G**) identified a number of endogenous chemicals including the fatty acids palmitic acid and palmitoleic acid; and cholesterol and cholesterol metabolites. This implicates a potential role of these organic compounds as zeitgebers important for maintaining circadian rhythms within the gut transcriptome. Additionally, SREBF (sterol regulatory element binding transcription factor) 1 and 2 were identified as transcriptional regulators of genes which lost rhythmicity (**Figure 3G**). SREBF1/2 are noted for their circadian rhythmicity in expression levels and activity, driven in part by feeding-fasting cycles^(36–38)^. Expression of *Srebf1* exhibited 24h rhythmicity in IECs under *ad libitum* feeding (peaking during the active phase) which was lost under interval feeding (**Supplementary Figure 4B**).

### Diurnal feeding behaviour is required for rhythmic IgA secretion

Intriguingly, amongst the pathways disrupted in IECs under interval feeding, we identified “Intestinal immune network for IgA production”. IgA plays a major role in host regulation of the microbiota, and its production by intestinal plasma cells is supported by IECs. Furthermore, we previously demonstrated that rhythms in IgA (generated in response to feeding) impacts upon the gut microbiota. Whilst some components of this pathway maintained 24h rhythms in IECs (*Itga4*, *Tnfsf13* and *Itgb7*), others lost (*Tgfb1*, *Ltbr*) or gained (*Il15ra*) rhythmicity (**Figure 3H**). Interestingly *Pigr* (encoding PIgR protein which facilitates IgA transport across epithelial cells) was highly expressed in IECs, but did not exhibit 24h rhythmicity under either feeding regimen. Altogether, these findings suggest that IECs may play further roles in facilitating the IgA response in a circadian manner. This prompted exploration of the impact of interval feeding on fecal IgA, which revealed a striking loss of 24h rhythms in IgA secretion (**Figure 3I**).

### Diurnal feeding is important for maintenance of rhythms in the microbiome

Given the importance of 24h rhythmicity in both IECs and IgA production for maintenance of the rhythmic microbiota^(9, 21, 39)^, we examined the response of the gut microbiota to interval feeding. 16s rRNA seq analysis was performed on fecal samples collected around the clock in *ad libitum* and interval fed mice. Analysis of observed richness (alpha diversity, Shannon index) revealed no difference between feeding regimen when all samples were pooled irrespective of collection time (**Supplementary Figure 5A**). However, as expected^(39)^ in *ad libitum* fed mice, richness exhibited variation over the 24h day (JTK_cycle analysis, adj. P<0.005), with a nadir during the rest phase. Although under interval feeding there was still a clear peak (ZT0) and trough (ZT8-12), the alpha diversity was not determined to be 24h rhythmic (JTK_cycle, P=0.23) (**Figure 4A**). Analysis of 24h rhythmicity in bacterial abundance in *ad libitum* fed mice revealed 231/477 oscillatory OTUs (total abundance, JTK_cycle analysis, adj. P<0.05), which was reduced to 103/477 under interval feeding (with an overlap of 60 OTUs) (**Figures 4B-D**). When OTUs were mapped onto phyla, *Bacteroidetes* and *Firmicutes* showed the expected anti-phasic relationship under both feeding conditions with interval feeding inducing a phase advance (**Figure 4E and Supplementary Figure 5B**). OTUs which lost rhythmicity included those assigned to *Lactobacillus gasseri* and *Oscillobacter* (**Figure 4F and Supplementary Figure 5C)**. Those that gained rhythmicity included *Clostridiales vadin BB60* group (**Figure 4G and Supplementary Figure 5C**) and those that remained the same included *Lachnospiraceae and Parabacteroides goldsteinii* (**Figure 4H and Supplementary Figure 5C**). Thus, these data highlight the importance of diurnal feeding behaviour on maintenance of a rhythmic microbiota. The microbiota plays a critical role in host metabolism, generating metabolic products from the breakdown of dietary components, with potent effects on the host. As an assessment of microbial outputs, SCFA levels were quantified across time from the caecum (**Figure 4I** and **Supplementary Table 1**). Of the 5 metabolites screened, caproic acid and valeric acid showed 24h rhythmicity (JTK_cycle, BH.Q<0.05) in *ad libitum* fed mice. However, these SCFAs lost rhythmicity in interval fed mice. Together these data demonstrate significant temporal re-organisation of the bacterial composition of the gut microbiota and its metabolic activity in the absence of diurnal feeding cues.

**Figure 4:**
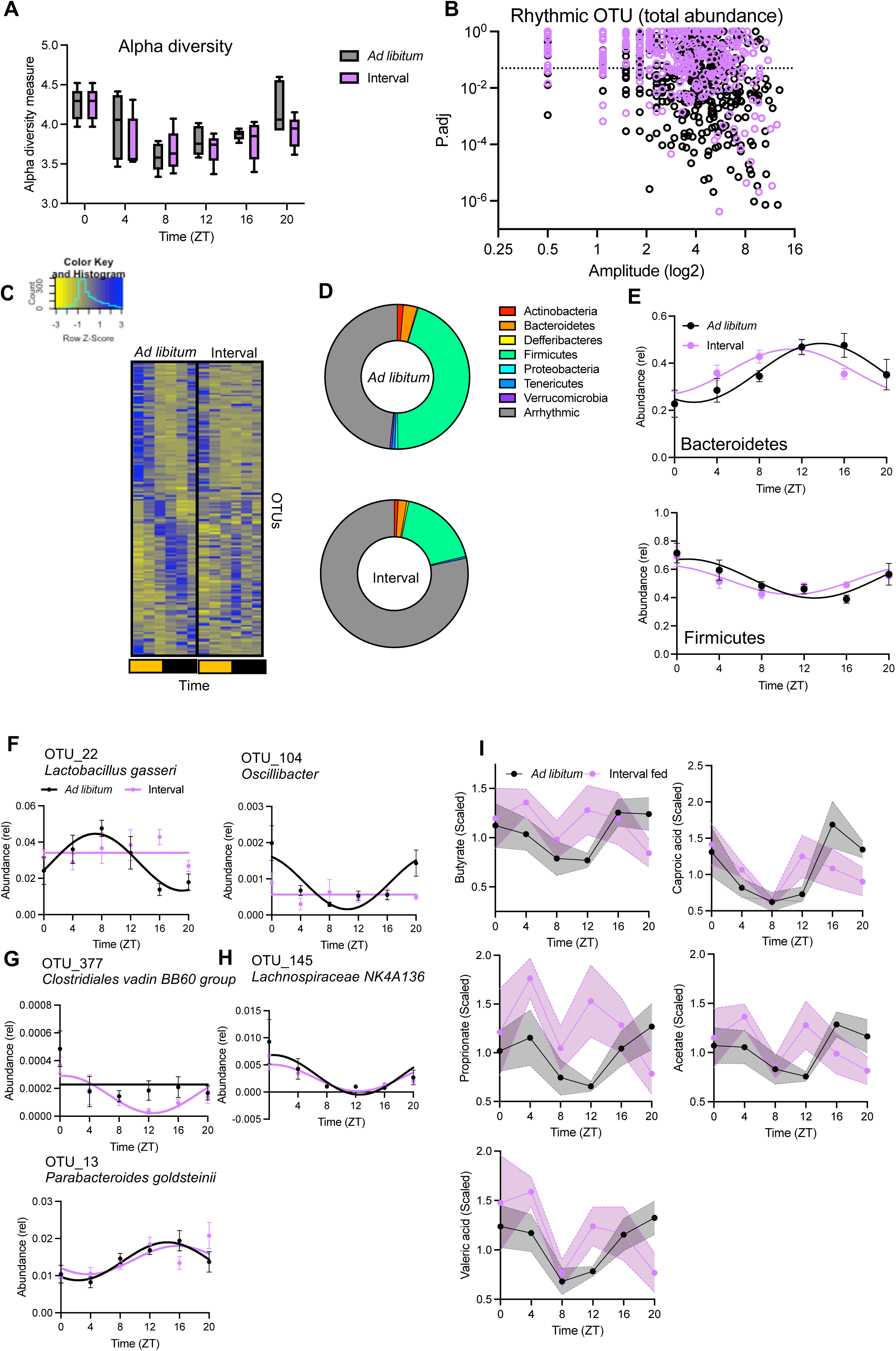
Interval feeding drives loss of rhythmicity in the microbiome. 16s rRNA seq analysis of fecal samples collected across the 24h day in *ad libitum* and interval fed mice. (A) Alpha diversity (Shannon Index) plotted across time, horizontal lines represent group mean and whiskers represent min to max, n=5/time point/feeding regimen. (B) Analysis of 24h rhythmicity (JTK_Cycle) of all OTUs (total abundance, 477 OTUs) from *ad libitum* (black) and interval (purple) fed mice. Points falling below the dotted line (P.adj<0.05) were assigned as rhythmic. (C) Relative abundance of OTUs (ordered by peak phase in *ad libitum* fed mice) where mean relative abundance is greater than 0.05%. (D) Taxonomic assignment by Phyla of rhythmic OTUs under the two feeding regimens. (E) Temporal distribution of OTUs (relative abundance) assigned to the two major phyla – Bacteroidetes and Firmicutes. Data was tested for fitting to a sine wave with a nonzero baseline and wavelength of 24. Example OTUs which (F) lost or (G) gained rhythmicity under interval feeding or were (H) unchanged. Data was tested for fitting to a sine wave with a nonzero baseline and wavelength of 24, and as appropriate a sine wave or horizontal line plotted. (I) Mass cytometric analysis of short chain fatty acids in caecal samples across time. Values are normalised to set metabolite median value to one, n=5/timepoint/condition. *See also Supplementary* Figure 5 *and Supplementary Table 1*.

## Discussion

This study demonstrates the importance of diurnal feeding behaviour on maintenance of 24h rhythmicity and host-microbiota interactions within the GI tract. The IECs house a robust molecular clock, which persists in the absence of diurnal feeding rhythms. Despite minimal disruption to the rhythmic nature of core clock genes, a significant proportion (∼25%) of the IEC circadian transcriptome lost rhythmicity under interval feeding, implicating reliance on feeding derived rhythmic zeitgebers, which may include fatty acids and sterols. 16s rRNA sequencing, IgA profiling and metabolomic analyses reinforces the notion that feeding behaviour regulates daily rhythms in secretory IgA and within the gut microbiota and its metabolic outputs. Together this work reveals that meal timing is a critical determinant of microbial rhythmicity and implicates the importance of feeding behaviour in maintenance of metabolic and immunological processes reliant on a rhythmic microbiota.

Mice with *ad libitum* access to food do not undergo significant periods of fasting *per se*, however their feeding behaviour is partitioned so that the majority (∼75%) of their daily food is consumed during the night. Interval feeding paradigms, sometimes termed “ultradian feeding” (where ultradian refers to a recurring rhythm with a period shorter than 24h) or “spread-out feeding”, have been used previously to abolish rhythmic food intake. Collectively, these studies have demonstrated impact on: peripheral gene expression over 24h^(30–32)^; the central clock^(33)^; rhythmic parasite behaviour^(34)^; and longevity driven by caloric restriction^(35)^. The interval feeding paradigm utilised here effectively partitioned food consumption into regular, equally sized meals, and rhythms in food intake were effectively uncoupled from behavioural rhythms. However, as a caveat, we note that overall daily food intake was somewhat reduced for interval fed mice, and these animals did not gain weight during the experimental period, whilst *ad libitum* fed mice did.

IECs are highly rhythmic with 36% of the transcriptome exhibiting 24h oscillations under *ad libitum* fed conditions. Through use of interval feeding, we demonstrated that diurnal rhythmicity in feeding behaviour is not required in order for the IEC clock to oscillate. This aligns with work demonstrating persistence of the hepatic molecular clock under a similar interval feeding regimen^(30)^. The IEC clock remained robustly rhythmic, however both the positive and negative arms of the clockwork machinery exhibited phase advances (with no change in amplitude). Several rhythmic feeding derived signals have demonstrated modulatory impact on the core clock. Amongst them are hormones such as insulin and insulin-like growth factor 1 (IGF-1)^(27)^; tryptophan metabolites^(28)^; and microbial metabolites such as SCFAs^(29, 40)^. Thus, whilst diurnal rhythms in feeding are not required to maintain IEC clocks, our data implicates the importance of rhythmic feeding derived cues for phase-setting and likely synchronisation of intrinsic clocks across multiple tissues. Although beyond the scope of this current study, a greater understanding of interactions between feeding derived rhythmic signals and the core clock machinery across peripheral tissues is warranted.

Because the IEC intrinsic clock persisted under interval feeding, we were able to interrogate the importance of autonomous clocks on the IEC transcriptome independent of the acute effects of feeding. Approximately a quarter of all cycling transcripts under *ad libitum* conditions lost rhythmicity in IECs in response to interval feeding. This suggests significant dependence on signals derived from rhythmic food intake. Thus, the IEC clock has only minor input to local transcriptional rhythmicity, and behavioural rhythms in feeding, driven by the central clock, provides a more significant contribution. It remains to be seen whether this dominant influence of feeding behaviour persists in other peripheral tissues beyond the gut and liver. Our analysis highlighted lipids (sterols and cholesterol) and fatty acids (palmitic acid and palmitoleic acid) as up-stream regulators of this subset of transcripts. Palmitate (an ester of palmitic acid) alters the circadian transcriptome via histone modification of enhancers^(30)^ and thus presents as a viable feeding derived signal important for driving rhythmicity in IECs independent of the clockwork machinery. Additionally, SREBF1/2 emerged as upstream transcriptional regulators of genes which lost rhythmicity. Expression and activity of these transcription factors exhibit robust 24h oscillations in the liver, driven largely by sterol availability^(36, 38)^, but also via the core clock component REV-ERBα^(41–44)^. Thus, rhythms in sterol availability provides a further feeding derived mechanism which could drive rhythmic transcription in the IEC independent of the core clockwork.

In the absence of diurnal feeding behaviour, we observed loss of rhythmicity within the microbiota. Prior work assessing the importance of feeding behaviour on the rhythmic microbiome has often relied on time restricted feeding paradigms^(19)^ or starvation^(39)^. Both approaches demonstrate significant impact, however the former is associated with inversion of the molecular clock within peripheral tissues^(45)^ (making it impossible to pick apart the relative contribution of feeding alone) and the latter is associated with metabolic stress. By uncoupling feeding behaviour and IEC clock activity, we reveal that feeding patterns have a profound effect on the rhythmicity of the gut bacteria, with interval feeding rendering 75% of the normally rhythmic OTUs arrhythmic.

The host immune system employs multiple mechanisms to orchestrate rhythms in the gut microbiota. This includes input from the IECs^(39)^, intestinal IgA^+^ plasma cells^(21)^, and innate lymphoid cells (ILCs)^(46)^. Given that interval feeding was not associated with disruption to the molecular clock in the IEC or liver, we speculate that intrinsic clocks within ILCs and plasma cells is similarly unaffected. We propose that interval feeding disrupts microbial oscillations via perturbation of rhythms in the availability of nutrients which influence IgA secretory activity of plasma cells. Our prior work has shown that feeding behaviour imprints rhythmicity onto the microbiome via rhythmic secretion of IgA by intestinal plasma cells, which is entrained by the timing of feeding and the resulting daily oscillations in nutrient availability^(21)^. One potential mechanism which warrants further exploration is availability of dietary cholesterol. Oxysterols, such as 25-hydrocycholesterol (25HC), regulate plasma cell IgA production. Availability of 25HC to plasma cells is controlled by IEC expression of cholesterol 25-hydroxylase (CH25H), which facilitates the generation of oxysterols from cholesterol^(47, 48)^. Interestingly, we previously demonstrated cholesterol biosynthesis pathways to be highly rhythmic in intestinal IgA+ plasma cells, while IgA secretion was determined by the time of feeding^(21)^. Thus, the striking loss of IgA rhythmicity observed under interval feeding could be driven by altered rhythmicity in cholesterol availability or altered cholesterol handling in IECs. This remains to be directly tested, and it is important that future work considers the small intestine, given that this region is the major site of IgA production.

In line with loss of 24h rhythmicity in IgA, 24h oscillations were lost in key commensals. This included the firmicutes, *Oscillobacter* and *Lactobacillus gasseri*. *Oscillobacter* rhythms were damped, with reduced relative abundance during the late night (ZT20-ZT0) in interval fed mice. *Oscillobacter* metabolise cholesterol, and in humans a high abundance of this microbe is associated with lower cholesterol levels^(49)^ suggesting that the observed damped rhythms might result in altered rhythms in cholesterol metabolism. The observed loss of rhythmicity in microbial populations in the absence of diurnal feeding-fasting behaviour has important implications for metabolic health. In mice, rhythmic microbial function modulates time-of-day differences in metabolite availability and uptake by the host, and is important for driving rhythmic blood glucose levels^(21)^. Further, in humans an arrhythmic microbiome is associated with increased risk of type 2 diabetes^(20)^. Microbiome rhythmicity also has important implications for maintenance of immunity. For example, a rhythmic microbiome is required for immune homeostasis in the gut^(6)^. Furthermore, rhythmic microbial metabolites imprint rhythmicity onto inflammatory disease^(22)^.

In summary, this study highlights the importance of food timing for maintenance of 24h rhythms in gut homeostasis. Feeding derived signals are important for maintenance of the circadian IEC transcriptome, independent of the function of the IEC intrinsic clock. Interval feeding resulted in loss of 24h rhythmicity in IgA secretion which was associated with loss of rhythmicity in a number of commensal bacteria and their metabolic outputs. Such loss of rhythmicity in the microbiome has been associated with aberrant metabolic and immune health. Together, these data highlight the importance of considering both “what we eat” and “when we eat” for maintenance of gut immune and metabolic health. Lifestyle factors including: chronotype (an individual’s natural preference to sleep and wake at certain times of the day); shift-work; and intermittent fasting all influence feeding behaviour, and the impact of this should be considered. Shift-work is associated with both metabolic disease and chronic inflammatory disease^(50–54)^. This is often linked to disrupted sleep timing and quality, however our study provides support for the contribution of disrupted feeding behaviour to ill health associated with shift work^(55)^. Modification of meal timing presents an attractive, cost-effective and accessible, behavioural intervention that may have benefits on health. Such interventions could have potential application in the management of adverse health consequences associated with shift-work or age-related changes in circadian function.

## Resource availability

Requests for further information and resources should be directed to and will be fulfilled by the lead contact, Julie Gibbs. This study did not generate new unique reagents. IEC RNAseq data from this study have been deposited at ArrayExpress with accession code E-MTAB-14883, and are publicly available as of the date of publication. 16s rRNA seq data reported in this paper will be shared by the lead contact upon reasonable request.

## Acknowledgements

We would like to thank Andy Hayes and the University of Manchester Genomics Core Technology Facility and Leo Zeef and the University of Manchester Bioinformatics Core Facility. We are grateful for the help received from Dr George Taylor and the University of Manchester Biological Mass Spectrometry Core Facility. 16s rRNA Sequencing was performed by the University of Liverpool Centre for Genomics Research. We would like to thank the University of Manchester Biological Sciences Facility staff for support with animal husbandry and maintenance, and the University of Manchester Flow Facility. Felicity Hunter was funded by an MRC doctoral training programme award. JEG is a Versus Arthritis Senior Fellow (22625). MRH is supported by a Wellcome Trust Sir Henry Dale Fellowship (105644/Z/14/Z).

## Author contributions

Conceptualisation, JEG, MRH and KJE; Methodology, FKH and JEG; Investigation: FKH, PD, ALM, SD, JC and JEG; Resources, JEG; Writing – original draft, FKH and JEG; Writing – review and editing, FKH, KJE, MRH, PD and JEG; Visualisation, FKH, PD and JEG; Supervision KJE, MRH and JEG; Project administration FKH and JEG; Funding acquisition, FKH, KJE, MRH and JEG.

## Supplementary files

Document S1: Supplementary Figures 1-5 and Supplementary Tables 1 and 2.

---

**Supplementary Figure 1:**

(A) Total daily food intake (g) per cage across feeding regimen, n=3 cages (*ad libitum*) and 5 cages (interval fed). 2-way repeated measures ANOVA and post-hoc Sidak’s multiple comparisons test. (B) Weight change across feeding regimen n=5/group, 2-way repeated measures ANOVA and post-hoc Sidak’s multiple comparison tests, all NS. (C) Animal weights across feeding regimen at the start (day 0) and end (day 17), n=5/group, 2-way ANOVA and post hoc Sidak’s multiple comparisons test. (D) VO2 (E) VCO2 and (F) Heat production under *ad libitum* feeding (black) and interval feeding (purple) as measured in Phenomaster cages. Line plots show group average (n=6 interval fed, n=5 *ad libitum* fed).

**Supplementary Figure 2:**

(A) Flow cytometric analysis of IECs (EpCAM^+^ CD45^-^) harvested from the colon, representative of n=3. (B) Transcripts demonstrating no change in rhythmicity (“same”) under interval feeding, n=4-5/timepoint, 6 time points per condition. (C) Mean expression levels (across all time points) of transcripts assigned same, change, gain or loss compared in *ad libitum* and interval fed samples, error bars show median and interquartile range, distributions compared by Welche’s t test, all ns. (D) Spline plots of normalized expression of all genes in the “bile secretion” (Mmu_04976, 50 genes) and ”Mineral absorption (Mmu_04978, 37 genes) pathways which appeared in our data, error bars represent 95% CIs around the mean. (E) Expression of clock genes within IECs (normalized expression) over time and their rhythmicity as assigned by CompareRhythms, n=4-5/time point.

**Supplementary Figure 3:**

Expression of core clock genes (*Per2, Bmal1* and *Rev-erbα*) over time within the liver of *ad libitum* and interval fed mice. Target gene expression was normalised to β-actin and expressed relative to ZT0 in *ad libitum* fed mice. Values are mean±SEM, n=3-5, 2 Way ANOVA, first level significance reported.

**Supplementary Figure 4:**

(A) Pathway analysis (mouse KEGG 2019) of transcripts which lost rhythmicity under interval feeding. Bars quantify fraction of the pathway represented, and dots represent statistical significance (dotted line marks P=0.05). (B) Expression of *Srepb1* across time in colonic IECs, 24h rhythmicity was lost in interval fed mice (CompareRhythms analysis), n=4- 5/time point.

**Supplementary Figure 5:**

(A) Violin plot showing alpha diversity (Shannon Index) between feeding regimens, solid horizontal lines represent group median and dashed horizontal lines represent quartiles, n=30, unpaired T test not significant. (B) Relative abundance by phyla across time, n=5/timepoint. (C) Example OTUs (total abundance). Data was tested for fitting to a sine wave with a nonzero baseline and wavelength of 24, and as appropriate a sine wave or horizontal line plotted.

## Methods

### Animals

All experimental procedures were performed in accordance with the UK Animals (Scientific Procedures) Act 1986, subject to local ethical review and approval from the University of Manchester Animals Welfare and Ethical Review Body (AWERB). Mice were housed in the Biological Services Facility (BSF) at the University of Manchester with regulated temperature (19-23°C) and humidity (45-65%). Mice were housed under a 12:12 light:dark cycle in light controllable cabinets, whereby zeitgeber time 0 (ZT0) refers to lights on, and ZT12 refers to lights off. 8-12 week old, male, C57BL/6J mice (Charles River Laboratories) were used and were co-housed. Mice were provided with *ad libitum* access to water. Standard chow was provided either *ad libitum* or via the interval feeding paradigm described below. Animal were provided with Sizzle-Nest and environmental enrichment in the form of shelters, gnawing sticks and cardboard tubes.

### Interval feeding

Food availability was restricted to a short feeding window every 3 h. Upon initiating the interval feeding regimen, during the 4 meals in the light phase (ZT2, ZT5, ZT8 and ZT11) food hoppers were made available for a 30 min window. For the 4 meals during the dark phase (ZT14, ZT17, ZT20 and ZT23), food hoppers were made available for a 10 min window. On the fifth day onwards (in response to observations that mice were consuming more food during the mealtime immediately after lights on, compared to later mealtimes during the light phase) the ZT2 meal was reduced to 20 min, and the dark phase meals were increased to 12.5 min to maintain equal light and dark food intake. Food was weighed after each feeding window and body weight was tracked. Food was delivered manually, or using a programmable automated system (TSE systems). Large intact food pellets were provided during meals, to avoid smaller fragments falling onto the cage floor, and cages were regularly checked to ensure there was no residual food on the floor.

### Analysis of metabolic parameters

Following a 2 week period of interval feeding (or for controls *ad libitum* feeding) in standard individual ventilated cages, mice were transferred to the PhenoMaster (TSE systems) for analysis of metabolic and behavioural parameters. Here, mice were singly housed within a light tight cabinet. All mice were provided with Sizzle-Nest and a shelter and had free access to drinking water. For interval fed animals, the food hopper was filled for the duration of the mealtimes and then emptied (ensuring no food had escaped onto the cage floor). Control mice had *ad libitum* access to food, but hoppers were agitated during the interval fed

animals’ mealtimes to ensure consistency. The PhenoMaster captured data every 2 mins including water consumption, activity (infrared activity frame), oxygen consumption and carbon dioxide production. Animals were allowed to acclimatise for 15h before data was collected for analysis.

### Colonic IEC isolation

Colonic IECs were extracted using established methods^(6)^. In brief, dissected colons were opened out longitudinally and sectioned into four pieces, washed in ice cold PBS and incubated in PBS containing 2% FBS on ice. After 20min, the tissue was transferred to HBSS containing 2mM EDTA and 1mM dithiothreitol and placed in a shaking incubator (180rpm, 37°C, 15min). Subsequently, samples were vigorously agitated for 30s to dissociate the epithelium from the basement membrane before being passed through a 70μm filter. The effluent was centrifuged (432xg, 4°C, 5min) and the pellet, containing IECs, re-suspended in 350μL RLT plus buffer (Qiagen) and stored at -20°C. RNA was extracted using the RNeasy Plus Mini Kit (Qiagen) according to the manufacturer’s instructions. For purity checks, isolated colonic cells were stained with PE labelled EpCAM (G8.8, eBioscience) and BV510 labelled CD45 (30-F11, Biolegend) 1:200 following a 20min Fc block (anti-mouse CD16/CD32 eBioscience, 1:100). Cells were washed with and then re- suspended in FACS buffer (PBS+5% FBS) before analysis on a BD LSR II.

### Liver RNA extraction

Liver tissue was homogenised in Trizol using Lysing Matrix D tubes and a BeadMill homogeniser (Fisher Scientific). RNA was extracted using chloroform, then precipitated using isopropanol. After washing in 70% ethanol, the RNA pellet was re-suspended in RNase free water.

### QPCR

RNA was quantified (NanoDrop, Thermo Fisher Scientific) and converted to cDNA using High Capacity RNA to cDNA kits (ThermoFisher). Taqman primers and probes (see **Supplementary Table 2**) were utilised, and plates were run on a QuantStudio 1 machine (Applied biosystems) using Takyon ROX probe 2X qPCR MasterMix dTTP blue (Eurogentic Ltd). β-actin was utilised as a housekeeping gene.

### RNA sequencing

RNA was extracted from IECs and quality was determined using a 2200 TapeStation (Agilent Technologies). Library preparation and sequencing was performed by the University of Manchester Genomic Technologies Core Facility. Libraries were generated using the TruSeq Stranded mRNA assay (Illumina) according to the manufacturer’s protocol. Multiplexed libraries were analysed by paired-end sequencing on a HiSeq 4000 instrument (76 + 76 cycles, plus indices), then de-multiplexed and converted using bcl2fastq software (v2.17.1.14, Illumina). Adaptors were removed and ends were trimmed using Trimmomatic (v0.36). Reads were mapped against the mouse genome (mm10/GRCm38) using STAR (v2.5.3). Reads were then counted, normalised and annotated in R using the Rsubread (v1.28.1), edgeR (v3.30.3) and biomaRt (v2.44.0) packages. One sample was excluded from further analysis from the *ad libitum* fed ZT20 group as it did not meet our quality control standards (leaving n=4 in this group). Differential expression analysis was run in RStudio using edgeR (v3.28.1). Genes were differentially expressed (DE) if the false discovery rate (FDR) was <0.05 and log2 fold change of <-2 or >2. Rhythmicity analysis was performed in RStudio using the CompareRhythms package^(56)^. Differentially rhythmic genes were identified by comparing the rhythmicity profiles between two treatment groups to classify genes either as ‘arrhythmic’ or as a ‘gain’, ‘loss’, ‘change’ or the ‘same’ rhythmicity.

### 16s rRNA Seq

Faecal pellets were collected from the terminal end of the colon (1-2 pellets/mouse), snap-frozen in liquid nitrogen and stored at -80°C. Bacterial DNA was extracted from 250mg sample using the Powersoil kit (Qiagen, Germany) according to the manufacturer’s instructions. DNA was eluted in the final step using a 50μl volume. DNA was quantified using a Nanodrop 2000 (Thermo Fisher Scientific) using a 260/280 purity threshold >1.8. and normalised across all samples. 20μL of DNA (1ng/μl) was sequenced using primers spanning the variable region 2 (V4) of the 16s rRNA gene^(8)^ by the University of Liverpool Centre for Genomic Research using the Illumina MiSeq v2 platform (Illumina) to generate 250bp paired end reads. Raw FastQ files were trimmed and reads <15 bp were discarded. The trimmed FastQ files were then processed using the 16S Amplicon Analysis pipeline in Centaurus Galaxy Server (University of Manchester Bioinformatics Core Facility), generating OTU tables. Briefly, reads were paired, OTUs were clustered using VSEARCH, and chimeric sequences were removed. The resulting OTU data was mapped onto taxonomies using the Silva reference database (v138) with a sequence homology threshold of >97%. The generated OTU biom table was subsequently analysed in R studio using the Phyloseq package (V1.38.0)^(57)^ which enabled the extraction of information on total and relative taxonomic abundances, as well as the assessment of alpha diversity, using the Shannon Index. Rhythmicity analysis of OTUs and taxonomies was performed using JTK_cycle^(58)^ of MetaCycle (v1.2.0), with period length fixed to 24 hours. OTUs were determined to be significantly rhythmic if the Adj.P < 0.05.

### Corticosterone ELISA

Terminal blood was collected in MiniCollect K3EDTA tubes (Greiner Bio-One) and stored on ice prior to centrifugation (3000xg, 4°C, 10min). Plasma was transferred to cryovials and flash frozen in liquid nitrogen. Plasma corticosterone levels were determined by ELISA (Enzo Life Sciences, Inc) following the manufacturer’s instructions. Absorbance values were determined using the Promega GloMax plate reader at 405nm.

### IgA ELISA

For assessment of IgA, faecal pellets were collected from the terminal end of the colon (1-2 pellets/mouse), snap-frozen in liquid nitrogen and stored at -80°C. Samples were prepared for analysis by homogenisation in PBS. Pellets were transferred to lysing matrix tubes (MP Biomedicals), re-suspended in PBS (100mg/ml) and homogenised using a Bead Mill 24 Homogenizer (Fisher Scientific). Samples were centrifuged (200xg, 1min) and the supernatant diluted (1mg/ml) before further centrifugation 3 times for 5 minutes at 200xg, 8000xg and 10,000xg, removing the pellet each time. The final supernatant was further diluted 1:4 in preparation for analysis. Faecal IgA concentration was determined by an ELISA (Bethyl laboratories) following the manufacturer’s instructions.

### SCFA analysis

Caecal contents were snap frozen and stored at -80°C prior to analysis. Samples were re- suspended 1:10 (based on weight of sample) in nuclease free water and transferred to lysing matrix D tubes (MP Biomedicals) for homogenisation using a BeadMill homogeniser. Homogenised samples were centrifuged (18,800xg, 5min, 4°C) and supernatant stored at - 80°C. To quantify levels of SCFAs, targeted metabolomics was performed using liquid chromatography-mass spectrometry (LC-MS; Mass Spectrometry Core Research Facility at the University of Manchester). LC-MS analysis was performed using a SCIEX Exion LC system consisting of two AD high pressure gradient pumps, vacuum degasser, solvent valve, AC column oven and AC autosampler, coupled to a SCIEX 7600 ZenoTOF Q-TOF mass spectrometer with TurboV optiflow ion source running a 50 μm ESI probe. The system was controlled by SCIEX OS v3.0. For each sample, an area ratio was calculated using the peak area of an internal standard (in order to correct for any batch effects) and this was normalised to the median value of all samples.

### Statistical analysis

Statistical tests were conducted in GraphPad Prism and are specified in the figure legends where appropriate. Throughout * denotes p < 0.05; ** p<0.01 and ***P<0.005.

