## Supplemental Files for "Meal timing is a critical factor for maintenance of gut homeostasis around the clock"

### Slide 1
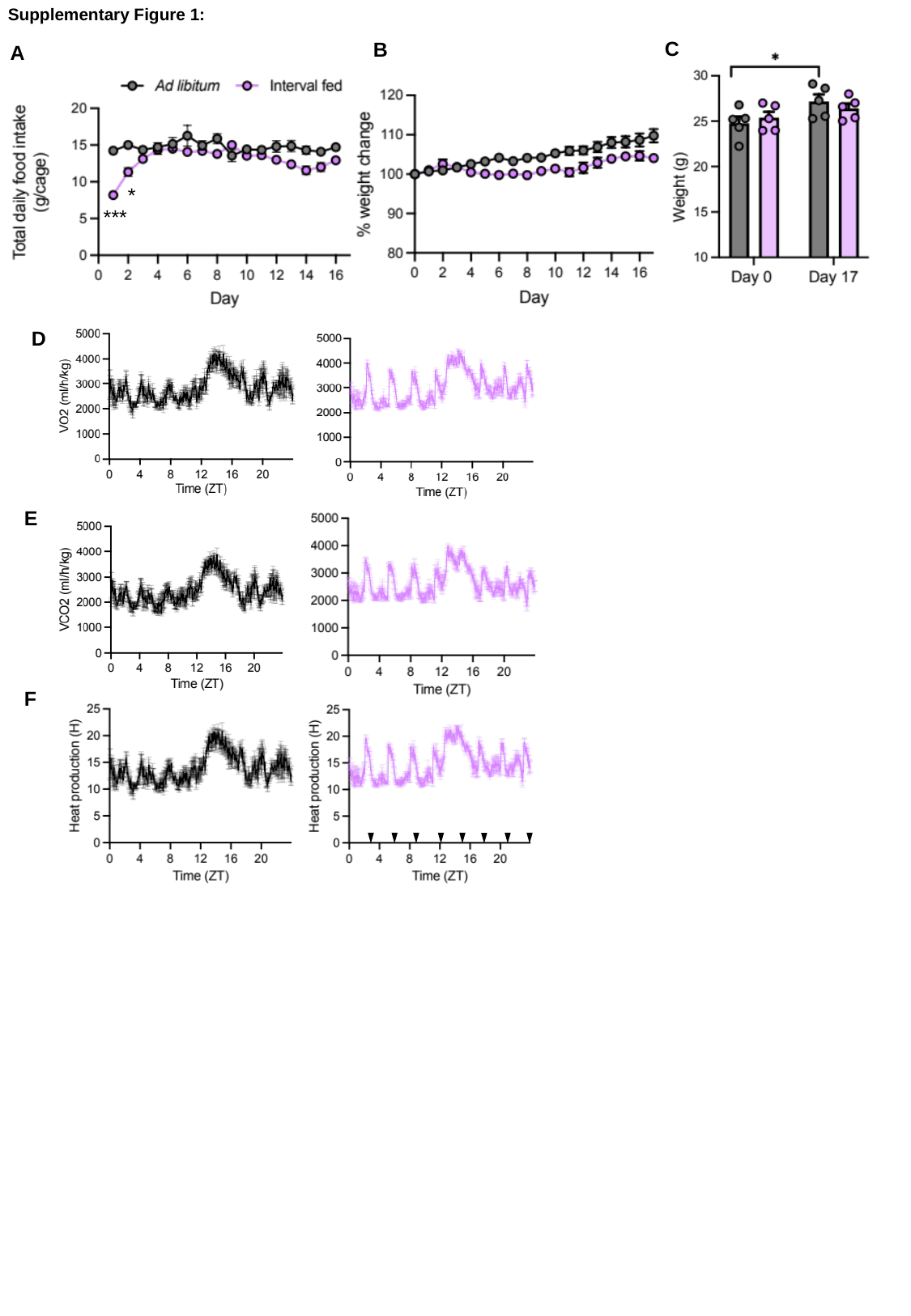

Supplementary Figure 1:
C
B
A
*
***
D
E
F

### Slide 2
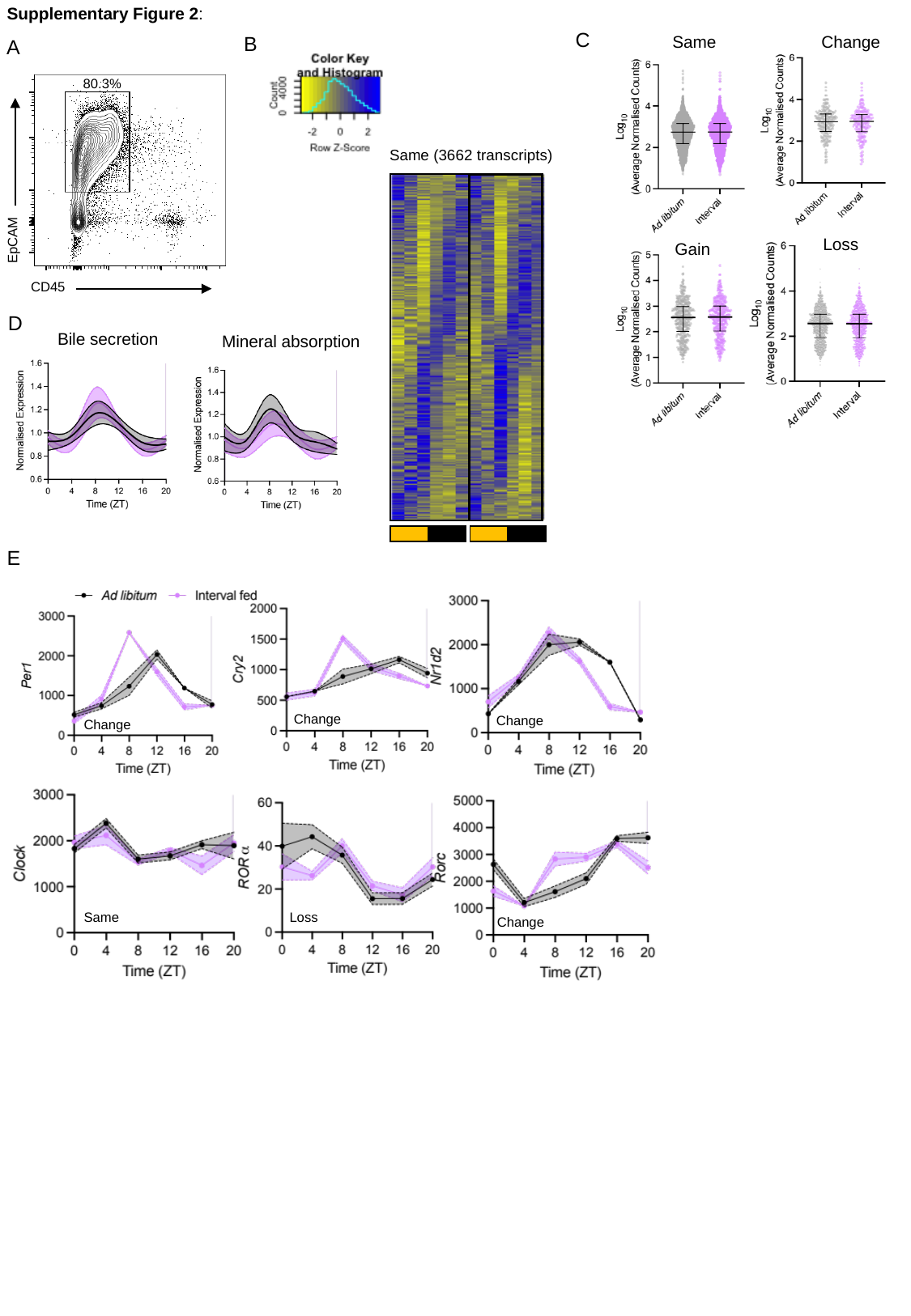

Supplementary Figure 2:
C
B
Same
Change
Gain
Loss
A
Same (3662 transcripts)
80.3%
EpCAM
CD45
D
Bile secretion
Mineral absorption
E
Change
Change
Change
Same
Loss
Change

### Slide 3
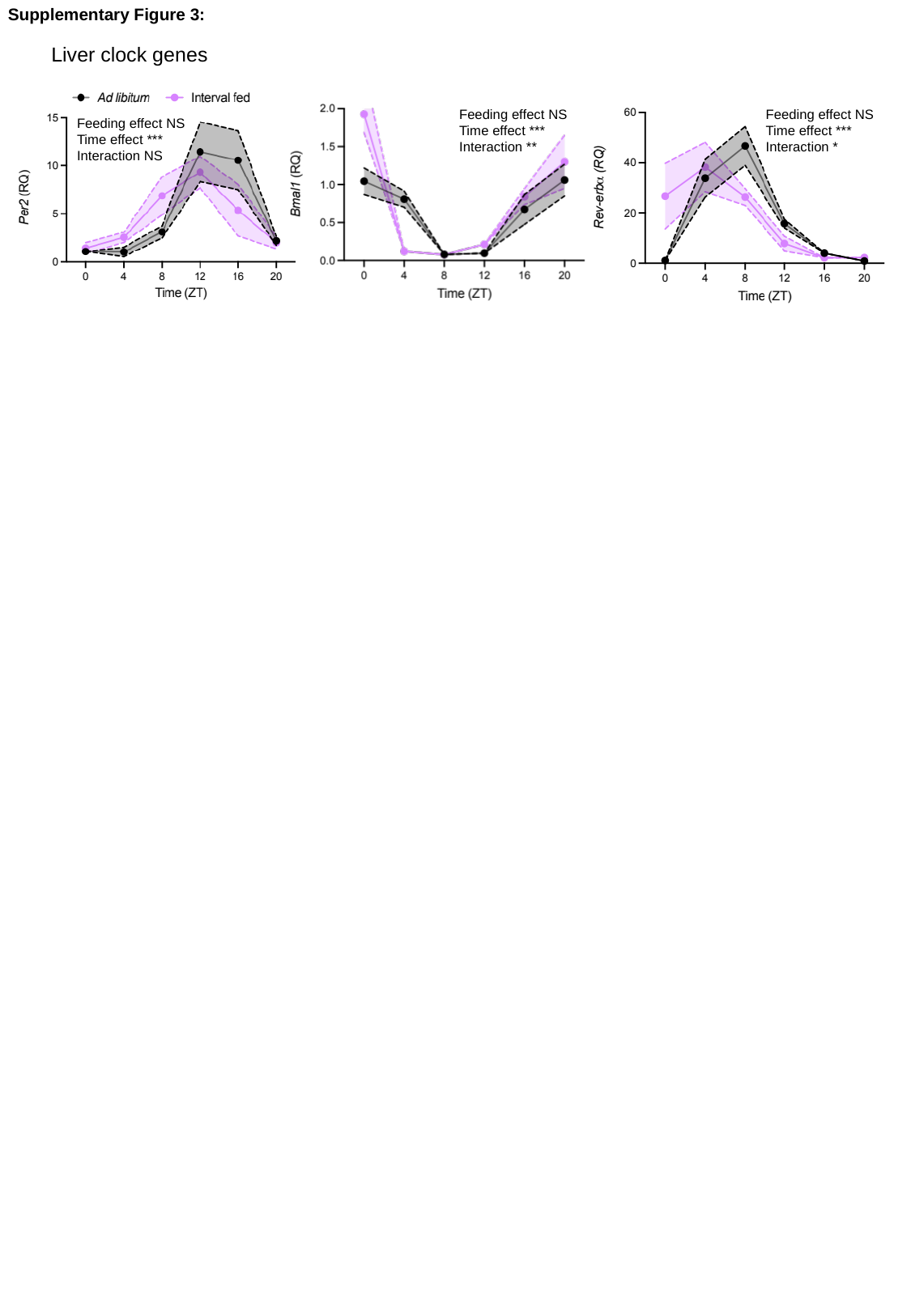

Supplementary Figure 3:
#### Liver clock genes
Feeding effect NS
Time effect ***
Interaction *
Feeding effect NS
Time effect ***
Interaction **
Feeding effect NS
Time effect ***
Interaction NS

### Slide 4
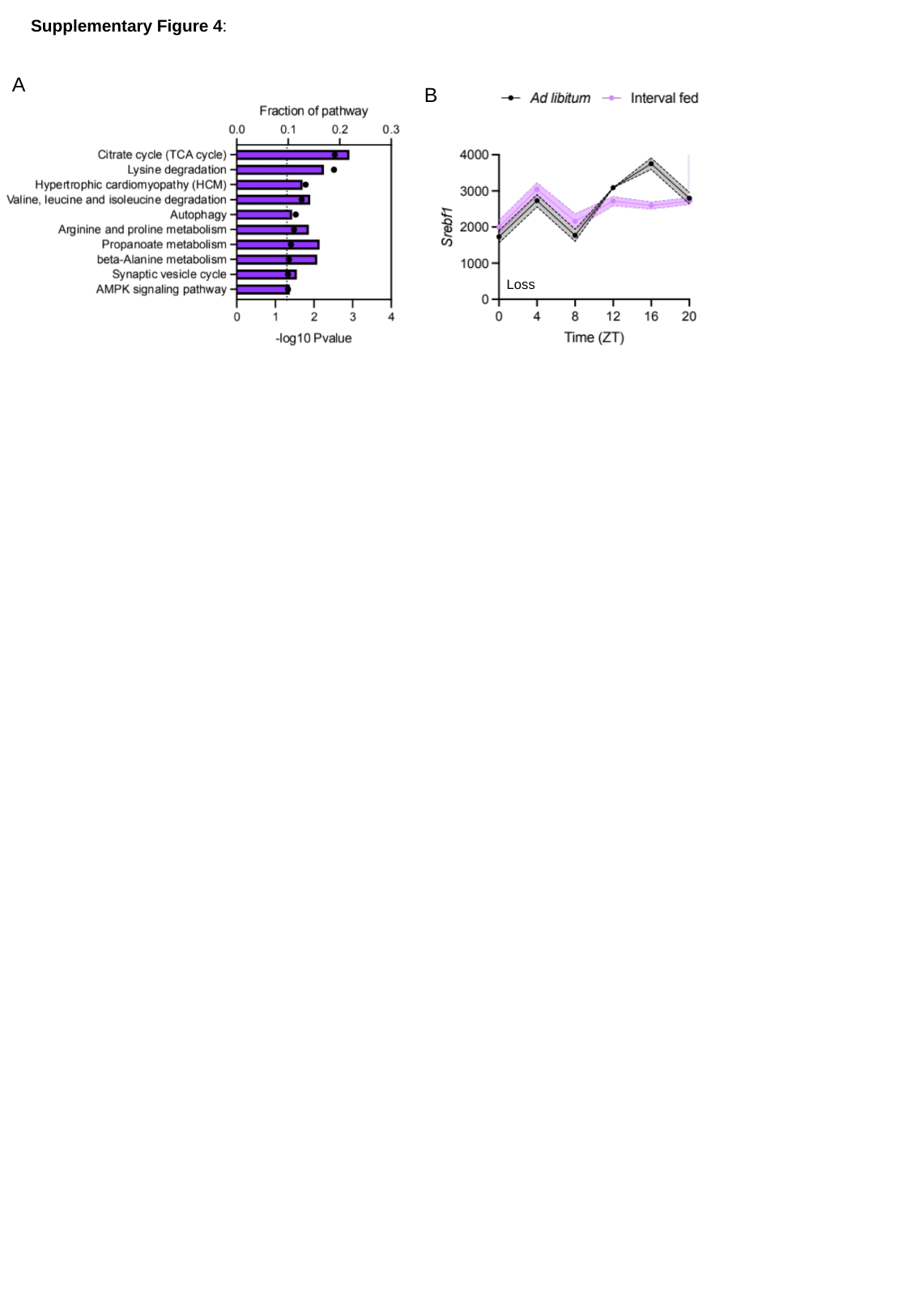

Supplementary Figure 4:
A
B
Loss

### Slide 5
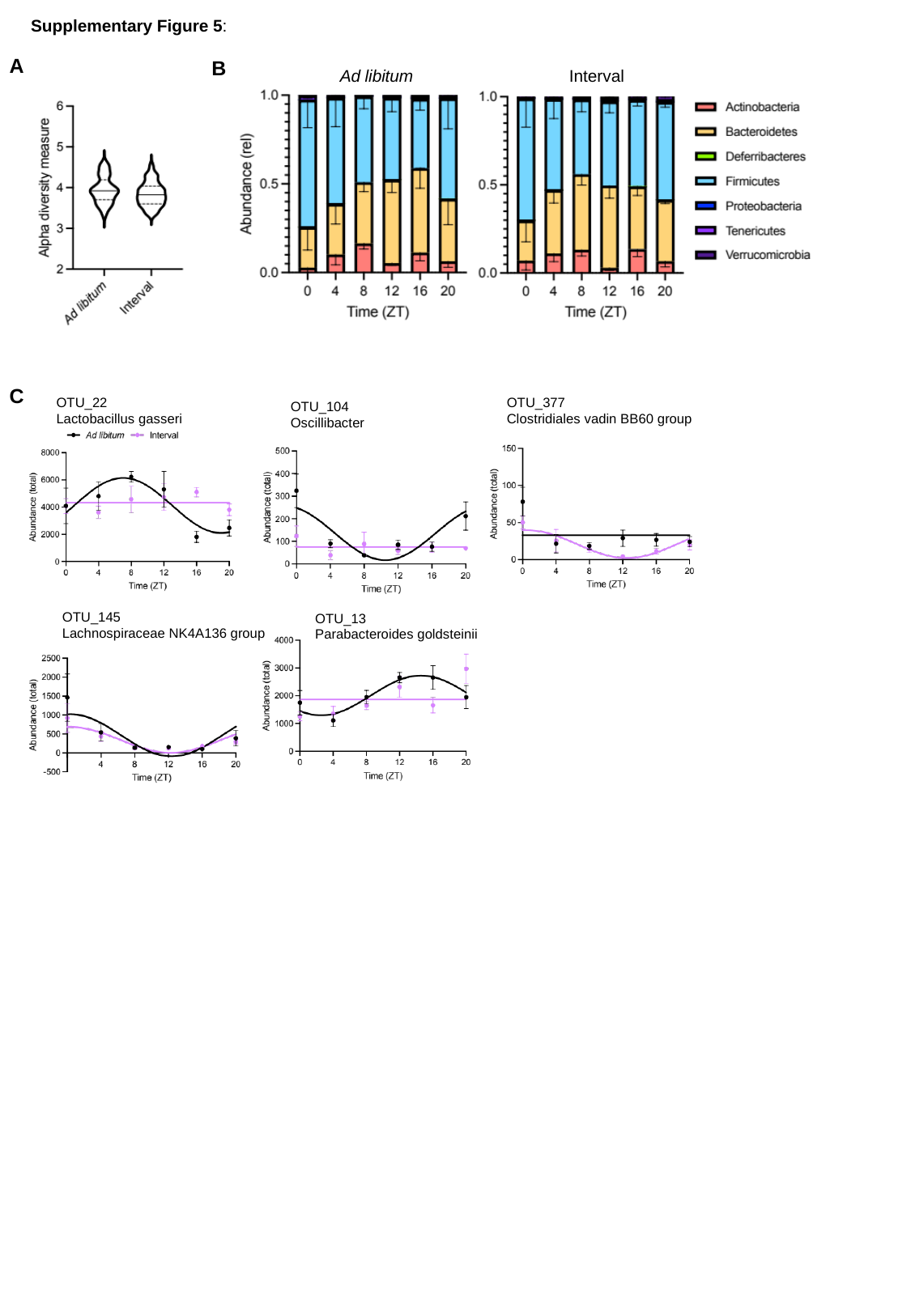

Supplementary Figure 5:
A
B
Interval
Ad libitum
C
OTU_22
Lactobacillus gasseri
OTU_377
Clostridiales vadin BB60 group
OTU_104
Oscillibacter
OTU_145
Lachnospiraceae NK4A136 group
OTU_13
Parabacteroides goldsteinii

### Slide 6
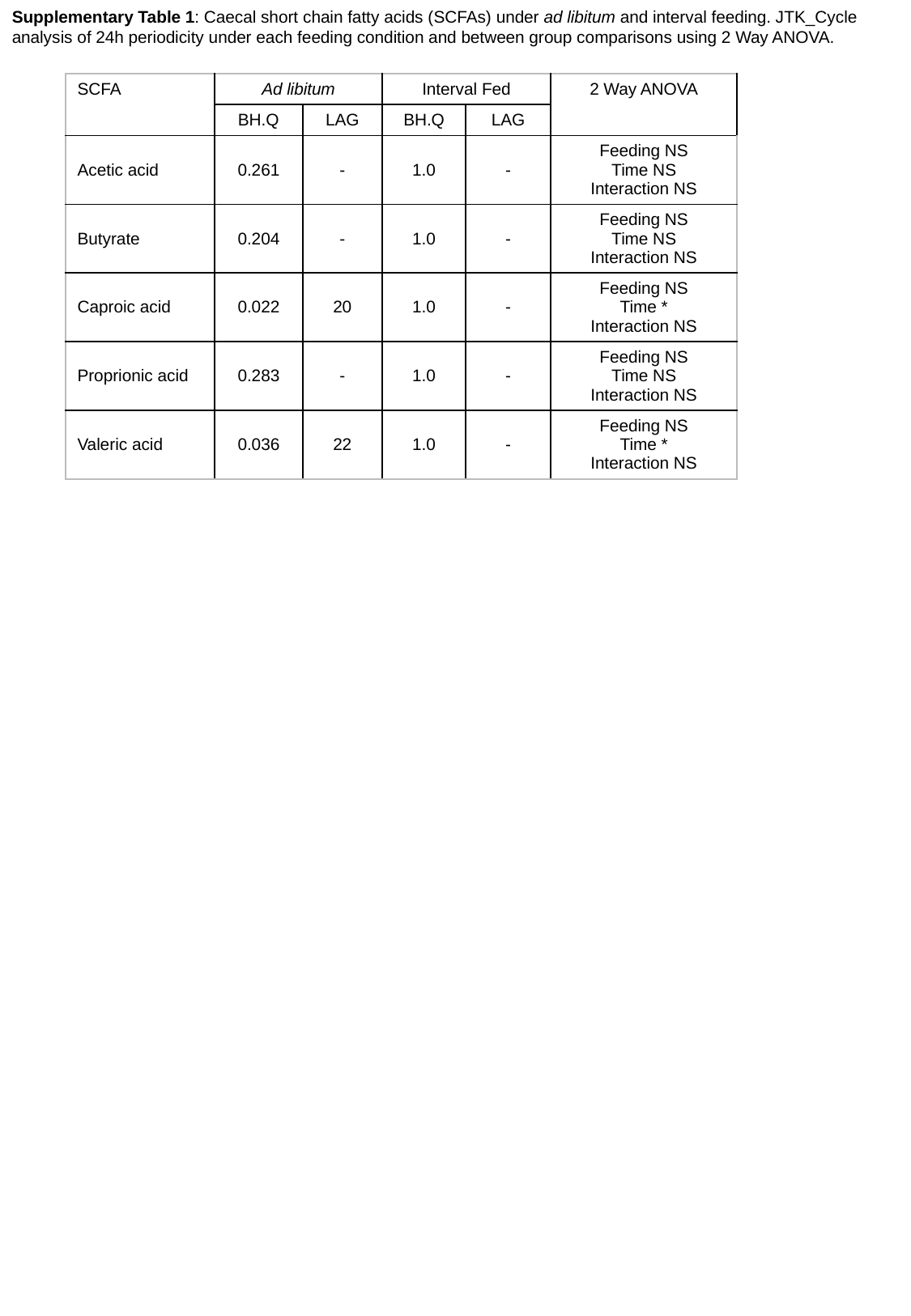

Supplementary Table 1: Caecal short chain fatty acids (SCFAs) under ad libitum and interval feeding. JTK_Cycle analysis of 24h periodicity under each feeding condition and between group comparisons using 2 Way ANOVA.
| SCFA | Ad libitum | | Interval Fed | | 2 Way ANOVA |
| --- | --- | --- | --- | --- | --- |
| | BH.Q | LAG | BH.Q | LAG | |
| Acetic acid | 0.261 | - | 1.0 | - | Feeding NS Time NS Interaction NS |
| Butyrate | 0.204 | - | 1.0 | - | Feeding NS Time NS Interaction NS |
| Caproic acid | 0.022 | 20 | 1.0 | - | Feeding NS Time \* Interaction NS |
| Proprionic acid | 0.283 | - | 1.0 | - | Feeding NS Time NS Interaction NS |
| Valeric acid | 0.036 | 22 | 1.0 | - | Feeding NS Time \* Interaction NS |

### Slide 7
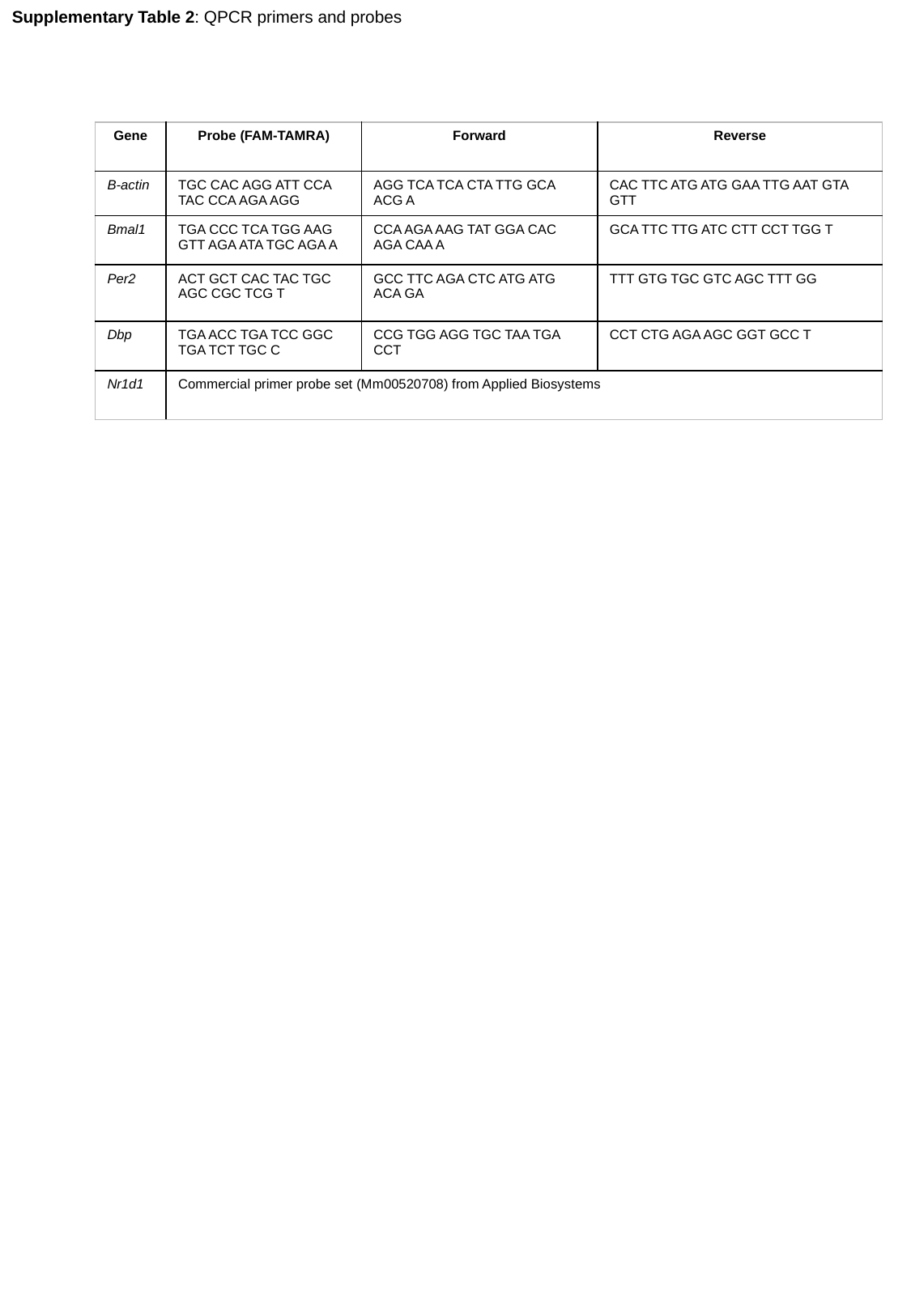

Supplementary Table 2: QPCR primers and probes
| Gene | Probe (FAM-TAMRA) | Forward | Reverse |
| --- | --- | --- | --- |
| B-actin | TGC CAC AGG ATT CCA TAC CCA AGA AGG | AGG TCA TCA CTA TTG GCA ACG A | CAC TTC ATG ATG GAA TTG AAT GTA GTT |
| Bmal1 | TGA CCC TCA TGG AAG GTT AGA ATA TGC AGA A | CCA AGA AAG TAT GGA CAC AGA CAA A | GCA TTC TTG ATC CTT CCT TGG T |
| Per2 | ACT GCT CAC TAC TGC AGC CGC TCG T | GCC TTC AGA CTC ATG ATG ACA GA | TTT GTG TGC GTC AGC TTT GG |
| Dbp | TGA ACC TGA TCC GGC TGA TCT TGC C | CCG TGG AGG TGC TAA TGA CCT | CCT CTG AGA AGC GGT GCC T |
| Nr1d1 | Commercial primer probe set (Mm00520708) from Applied Biosystems | | |
